# Suppression of miR-199a-5p alleviates ulcerative colitis by upregulating endoplasmic reticulum stress component XBP1

**DOI:** 10.1101/2021.02.05.430002

**Authors:** Shanshan Wang, Lei Shen, Shuai Peng, Minxiu Tian, Xiangjie Li, Hesheng Luo

## Abstract

Aims: This study aimed to explore the biological activities of miR-199a-5p in dextran sulphate sodium (DSS)-induced ulcerative colitis and apoptosis and identify the direct target of miR-199a-5p in this process. Main methods: HT-29 cells and C57BL/6 mice were used to examine the function of miR-199a-5p in vitro and in vivo, respectively. Expression of miRNA and mRNA was measured using quantitative real-time PCR and western blotting was used to measure the change in protein expression. Flow cytometry was subsequently employed to determine cell apoptosis, and a luciferase assay was used to confirm the direct target of miR-199a-5p. Results: Expression of miR-199a-5p was increased by DSS treatment in mice. In parallel, miR-199a-5p is found to be involved in endoplasmic reticulum stress (ERS) and cell apoptosis in HT-29 cells, and its upregulation induced ERS, apoptosis, weight loss, and ulcerative colitis in mice in vivo, which could be prevented by the suppression of miR-199a-5p. Luciferase assay confirmed that the 3′ untranslated region (3′-UTR) of XBP1 is the target binding site of miR-199a-5p. Conclusion: miR-199a-5p promotes ulcerative colitis and cell apoptosis by targeting the 3′-UTR of XBP1. Our findings reveal a new regulatory mechanism for ERS signaling and suggest that miR-199a-5p might be a potential target for UC therapy.

## Introduction

Ulcerative colitis (UC) is a nonspecific intestinal inflammatory disease with unclear etiology [1]. It is a chronic disease, the pathological processes of which are mainly recurrent intestinal inflammation, mucosal barrier injury, prolonged ulceration, and inflammatory hyperplasia. In addition, UC patients have a higher risk of colorectal cancer [2]. Intestinal infection, immunity, heredity, and other factors may lead to abnormal inflammatory processes in colonic mucosa. Endoplasmic reticulum stress (ERS) is thought to be one of the mechanisms leading to the development and progression of intestinal inflammation during the pathogenesis of UC [3], as it can activate the expression of protective proteins such as endoplasmic reticulum chaperones and enhance the ability of cells to resist stress. However, sustained ERS can cause an increase in unfolded proteins, leading to cell dysfunction and the transformation of ERS from a protective effect promoting cell survival to a pro-apoptotic one. In recent years, various studies have found that ERS is involved in the pathological process of UC by inducing cell apoptosis, participating in the innate immune response and epithelial autophagy [4, 5].

MicroRNA (miRNA) is a novel gene regulator that negatively affects the expression of target genes at the post-transcriptional level by directing the degradation of mRNA or inhibiting its translation. Bartoszewski first found the regulatory relationship between miRNA and ERS in 2011 [6]. After that, numerous studies have shown that miRNAs play an important role in the regulation of genes in ERS-related signal transduction pathways and protein translation. The mutual regulatory relationship between miRNA and ERS-related factors is an important reason for determining whether ERS is protective or damaging to cells.

Through previous bioinformatics analysis, we have predicted that miR-199a-5p mediatesd the ERS process by inhibiting XBP1 in the regulation of UC’s pathophysiological process. This study demonstrated this regulatory mechanism.

## Results

### Expression differences of miR-199a-5p and XBP1 in DSS-induced colitis mice

After treatment with 3% DSS for seven days, the body weight of the mice was greatly reduced compared with those given normal drinking water (Figure 1A), and their DAI score was higher than that of the control group (Figure 1B). The colon tissue structure of the control group was generally normal, and there was no obvious inflammatory cell infiltration in the lamina propria and submucosa. In the DSS group, the colonic mucosa of the mice was incomplete, glandular arrangement was disordered or even absent, and the inflammatory cell infiltration was obvious (Figure 1C). Similarly, miR-199a-5p expression was markedly increased in DSS-treated mice compared with mice given normal drinking water (Figure 1D) and the corresponding XBP1 expression decreased considerably (Figure 1E, F).

**Figure 1:**
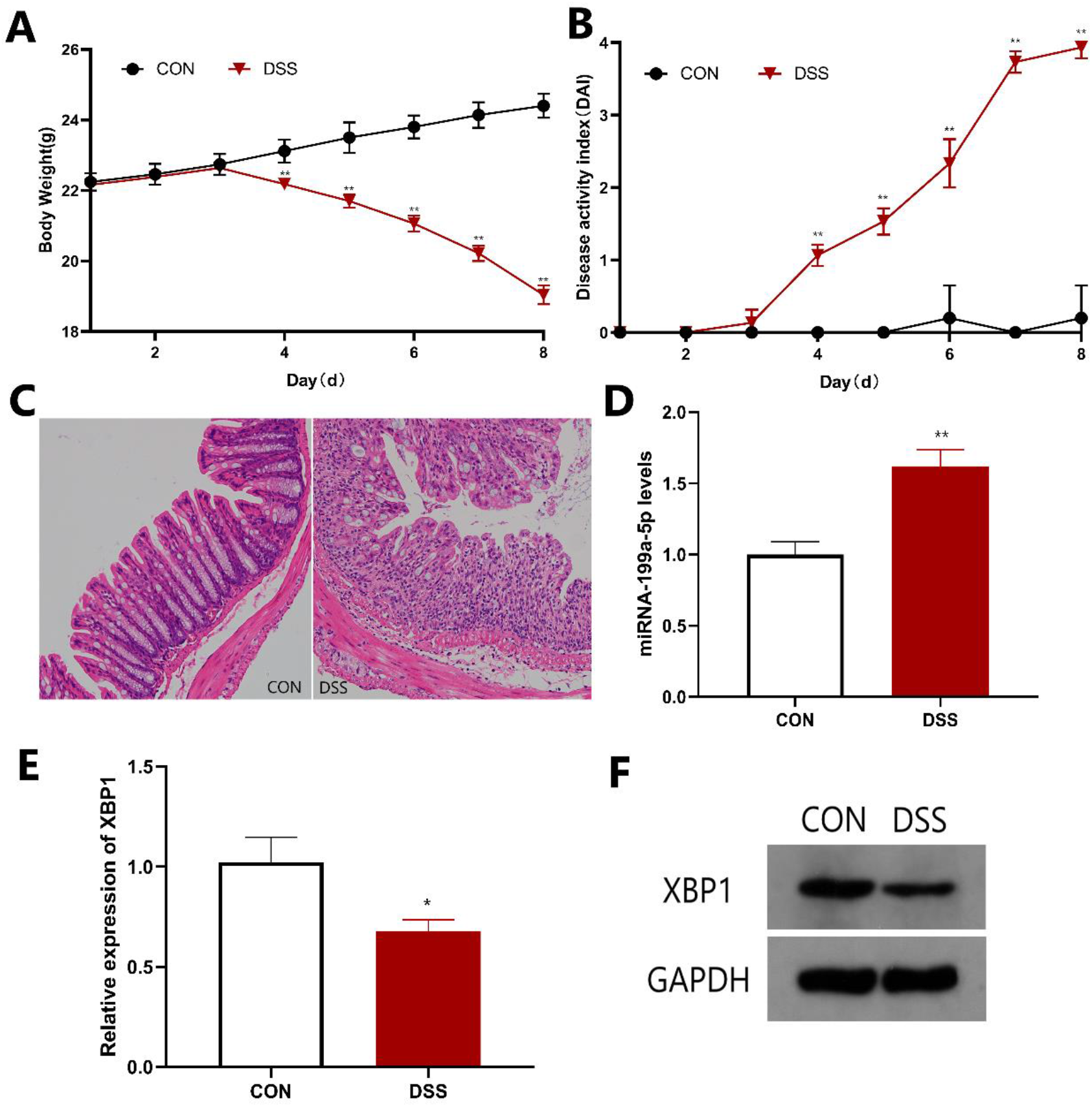
Expression differences of miR-199a-5p and XBP1 in DSS-induced colitis mice. A: Body weight was reduced in mice treated with DSS; B: DAI score was elevated in mice treated with DSS; C: Mucosal histology was examined by HE staining (200× magnification); D: miR-199a-5p expression was upregulated in mice treated with DSS; E, F: XBP1 expression was downregulated in mice treated with DSS. \**P* < 0.05 \*\**P*< 0.01.

### miR-199a-5p induces ERS in TG-treated HT-29 cells

miR-199a-5p expression was significantly upregulated in HT-29 cells treated with TG for 24 h (Figure 2A), indicating that the ERS process could lead to an increased expression of miR-199a-5p. After the introduction of miR-199a-5p mimics and inhibitors, TG was used to treat HT-29 cells. After eight hours, mRNA levels of ERS marker proteins GRP78 and XBP1 were significantly increased, with the most being in the inhibitor group. After 24 hours of treatment, GRP78 and XBP1 mRNA levels in the TG group and the mimic group decreased to less than those in the control group, while those in the inhibitor group remained higher (Figure 2B). The increase of GRP78 in the early stage of ERS indicates that the ERS self-repair process is initiated, explaining the increase in expression levels in each group at 8-h. GRP78 decreased with the extension of ERS process, indicating that the stress response of the endoplasmic reticulum decreased and cell function was impaired. The expression of GRP78 and XBP1 were detected by western blotting (Figure 2C), and compared with the TG-treated group, expression of both proteins in the inhibitor group was higher, indicating that inhibition of miR-199a-5p could alleviate intracellular ERS.

**Figure 2:**
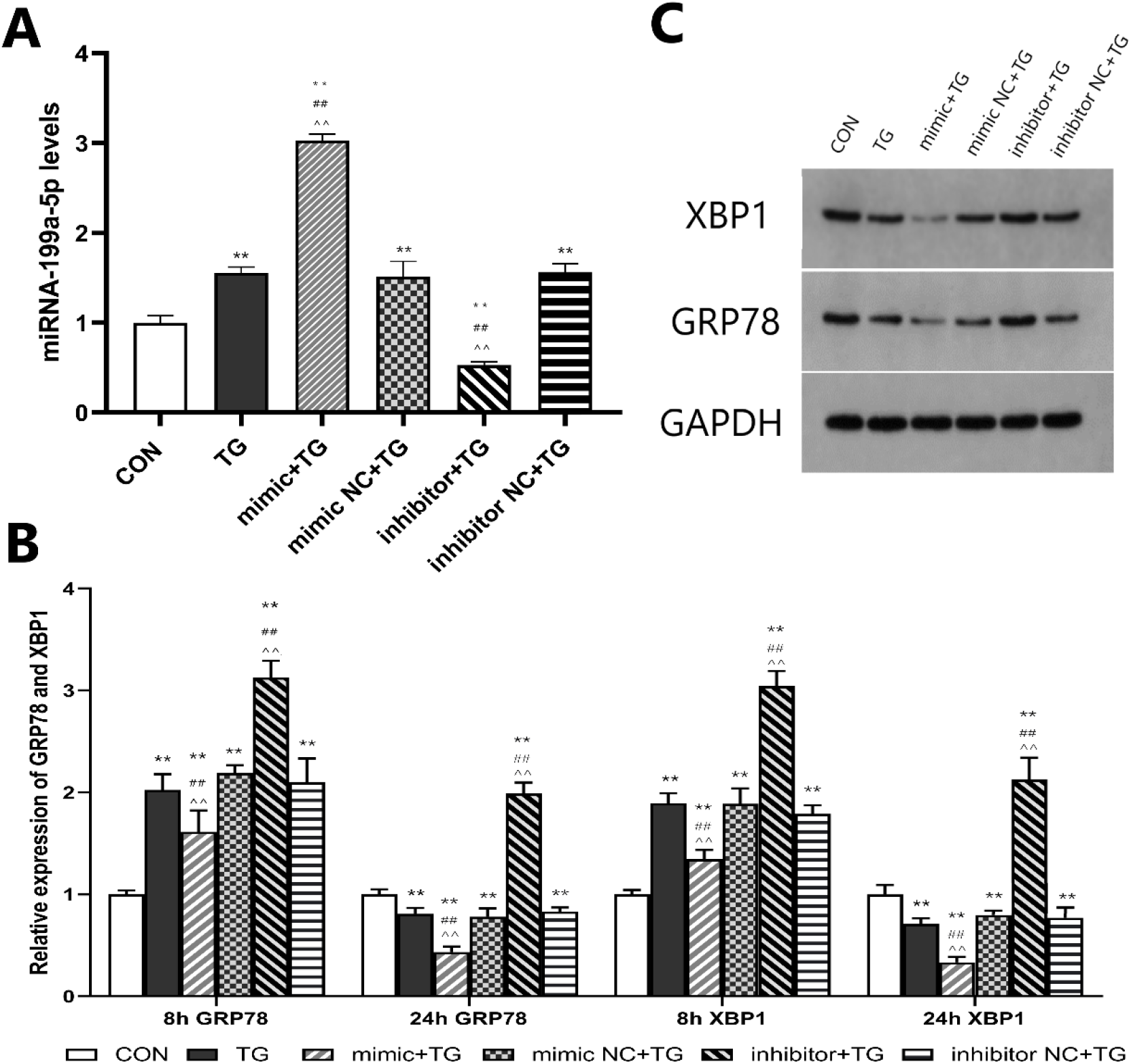
miR-199a-5p induced ERS in TG treated HT-29 cells. A: After transfection with miR-199a-5p mimics and inhibitors, HT-29 was treated with TG for 24 hours to detect the expression of miR-199a-5p; B: After transfection with miR-199a-5p mimic and inhibitor, HT-29 was treated with TG to detect the expression levels of GRP78 and XBP1 mRNA at 8 and 24 h, respectively; C: After transfection with miR-199a-5p mimics and inhibitors, HT-29 was treated with TG for 24 hours to detect the expression of GRP78 and XBP1. \*\**P*< 0.01 vs. CON; ##*P*< 0.01 vs. TG; ^^*P*< 0.01 vs. NC

### miR-199a-5p induces cell apoptosis in TG-treated HT-29 cells

Cell apoptosis was increased in TG-treated cells, and this increase was inhibited in groups with miR-199a-5p inhibitor (Figure 3A, B). The expression of the ERS and apoptosis-related proteins caspase-12 and CHOP was upregulated by TG, and the expression of the two proteins in cells also transfected with miR-199a-5p was lower than that in the other groups, suggesting that miR-199a-5p inhibition can alleviate cell apoptosis (Figure 3C).

**Figure 3:**
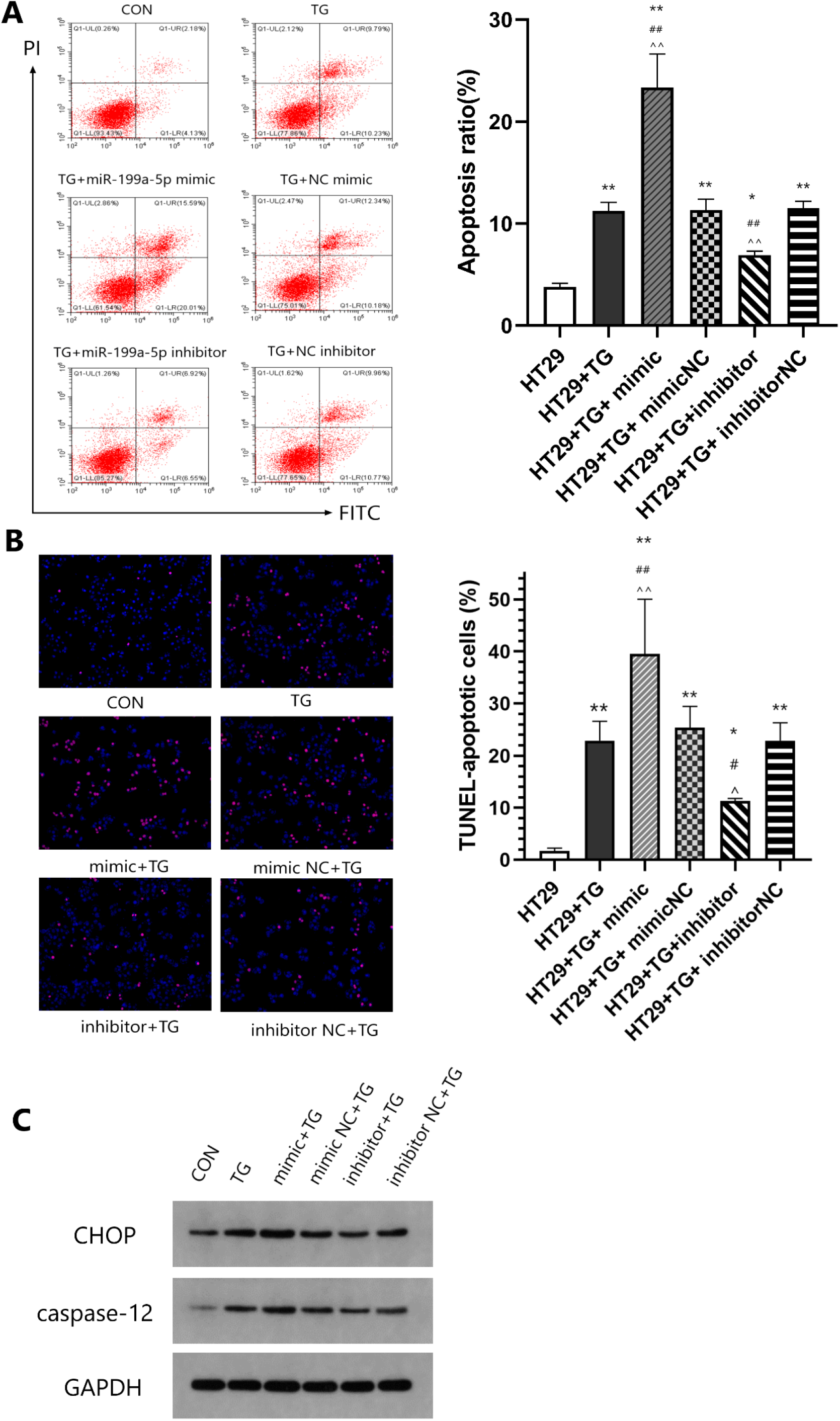
miR-199a-5p induced cell apoptosis in TG-treated HT-29 cells. A: Flow cytometry was used to detect apoptosis in each group; B: TUNEL detected apoptosis in each group; C: The expression of ERS-associated apoptotic proteins CHOP and caspase-12 in each group. miR-199a-5p inhibitor inhibited TG-induced cell apoptosis in HT-29 cells. \**P*< 0.05 \*\**P*< 0.01 vs. CON; #*P* < 0.05 ##*P* < 0.01 vs. TG; ^*P* < 0.05 ^^*P* < 0.01 vs. NC

### XBP1 is the direct target of miR-199a-5p

TargetScan was used to predict the potential target of miR-199a-5p. The results demonstrated that there was a miR-199a-5p binding site in the 3′-UTR of XBP1 (Fig. 4A). Co-transfection of XBP1 3′-UTR and miR-199a-5p mimics showed a reduction of luciferase activity compared to the negative control, while no significant difference was observed in cells transfected with XBP1 3′-UTR scramble sequence and miR-199a-5p mimic (Fig. 4B). Protein expression of XBP1 was reduced by the miR-199a-5p mimic and increased by the miR-199a-5p inhibitor (Fig. 4C, D).

**Figure 4:**
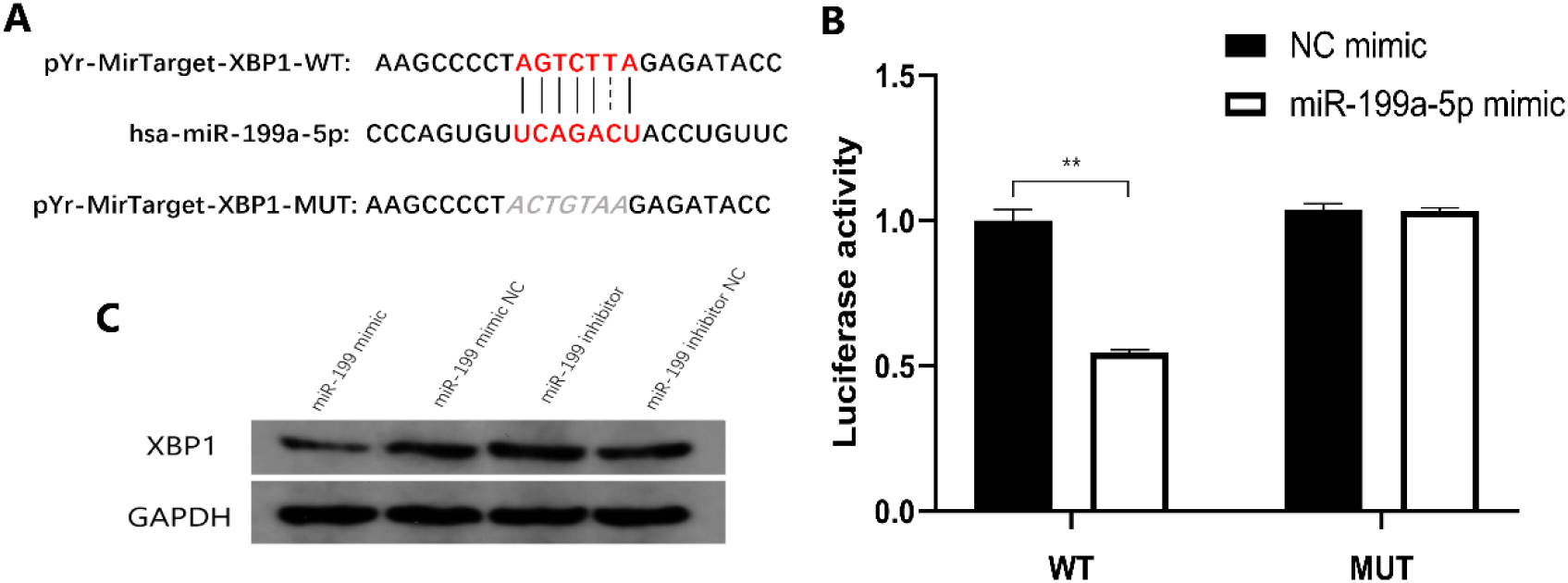
3′-UTR of XBP1 was the direct target of miR-199a-5p. A: The putative binding sites between miR-199a-5p and 3′-UTR of XBP1; B: Luciferase activity was decreased in co-transfection of pYr-MirTarget-XBP1 3′-UTR-WT and miR-199a-5p mimic, but there was no change in co-transfection of pYr-MirTarget-XBP1 3′-UTR-MUT and miR-199a-5p mimic; C: Protein expression was reduced by miR-199a-5p mimic and induced by miR-199a-5p inhibitor. \*\**P*< 0.01

### Inhibition of miR-199a-5p alleviates DSS-induced ulcerative colitis

Inhibition of miR-199a-5p was achieved by enema of rAAV-inhibited miR-199a-5p into mice. The body weight was reduced in mice treated with DSS, which was likely prevented through inhibition by miR-199a-5p (Figure 5A). DAI score increased in mice treated with DSS, and this increase was smaller in mice treated with rAAV-inhibited miR-199a-5p (Figure 5B). The colon of mice in the control group was normal, without obvious congestion and swelling. Compared with the control, the colon length of mice in the DSS group was significantly shortened, while the colon condition of mice in the inhibition group improved (Figure 5C). According to HE staining results, the pathological condition of the colonic mucosa in the inhibition group was relatively mild (Figure 5D). Upregulation of miR-199a-5p was observed in the DSS group, and transfection of rAAV-inserted miR-199a-5p reduced the expression of miR-199a-5p (Figure 6A). The expression of XBP1 decreased in DSS-treated mice. Compared with the control group, XBP1 expression was significantly upregulated in the inhibition group (Figure 6B). The expression of inflammatory cytokines IL-6 and TNF-α in colon tissues of mice in the DSS group was significantly increased, while it was decreased in the inhibition group (Figure 6C). The expressions of GRP78, XBP1, caspase-12, and CHOP were upregulated in mice treated by DSS, and GRP78 and XBP1 expressions increased when miR-199a-5p was inhibited, indicating that ERS was relieved in mice. Caspase-12 and CHOP decreased when miR-199a-5p was inhibited, suggesting that miR-199a-5p inhibition could alleviate apoptosis in DSS-induced mice (Figure 6D).

**Figure 5:**
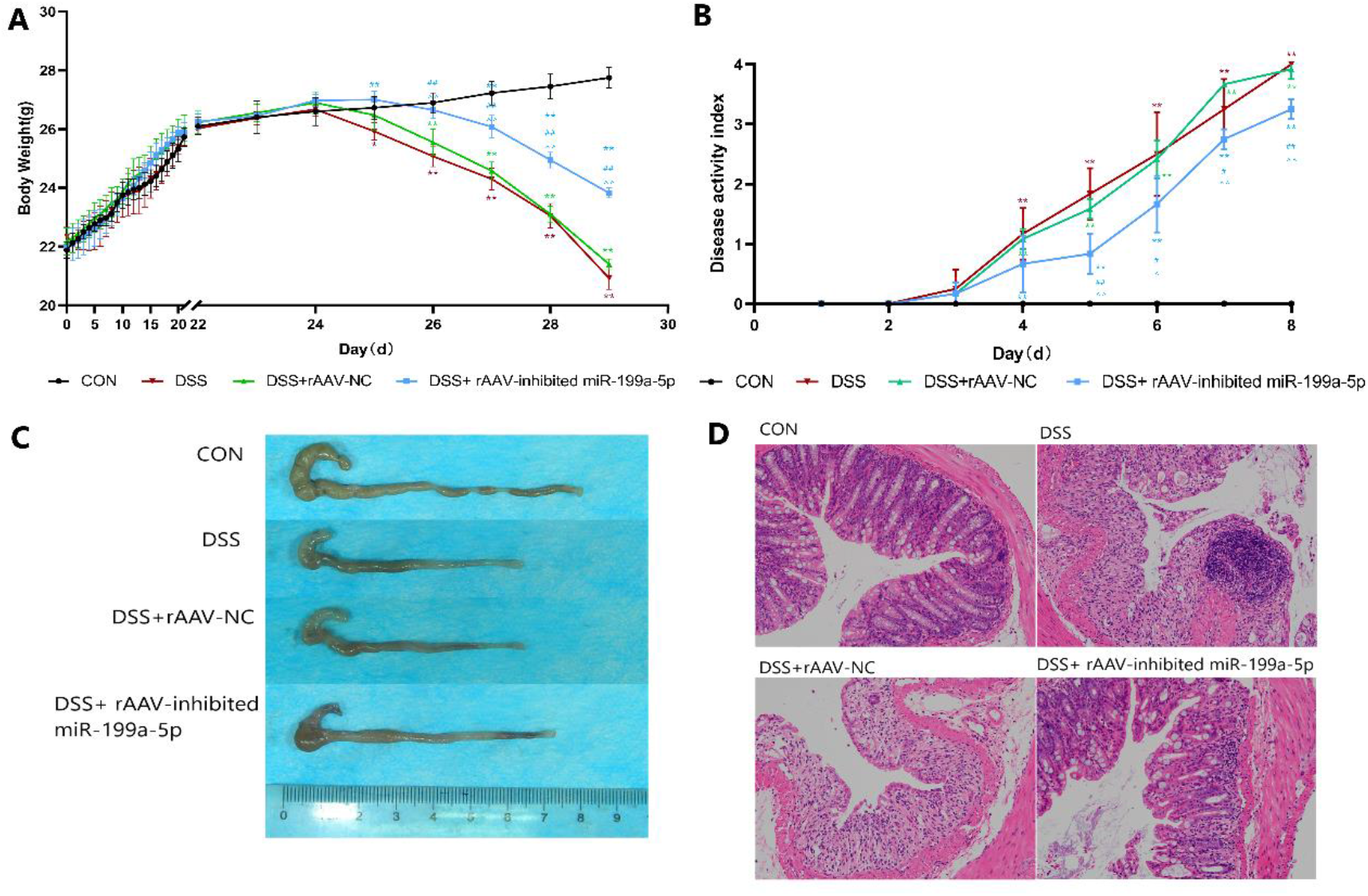
Inhibition of miR-199a-5p alleviated DSS-induced ulcerative colitis. A: Body weight loss induced by DSS was prevented by inhibited miR-199a-5p; B: Elevation of DAI score induced by DSS was decreased by inhibition of miR-199a-5p; C: Colon length of mice in each group; D: Mucosal histology was examined by H&E staining (200× magnification). \**P*< 0.05 \*\**P*< 0.01 vs. CON; #*P*< 0.05 ##*P*< 0.05 vs. DSS; ^*P*< 0.05 ^^*P*< 0.01 vs. NC

**Figure 6:**
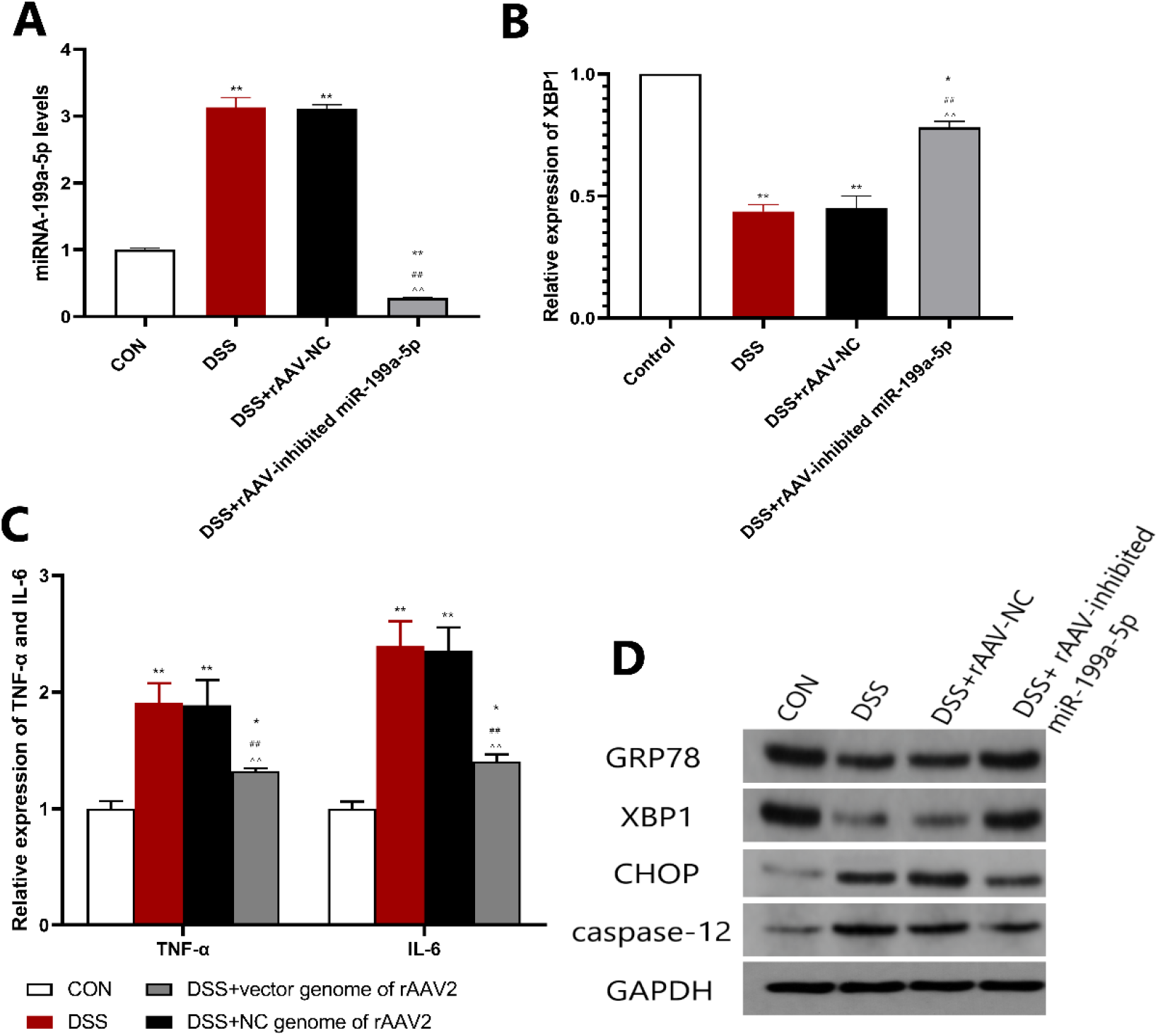
Inhibition of miR-199a-5p alleviated DSS-induced ulcerative colitis. A: Upregulation of miR-199a-5p induced by DSS was inhibited by rAAV-inhibited miR-199a-5p; B: Downregulation of XBP1 mRNA induced by DSS was inhibited by inhibited miR-199a-5p; C: Upregulation of IL-6 and TNF-α induced by DSS were inhibited by inhibited miR-199a-5p; D: Change in GRP78 and XBP1 expression induced by DSS was upregulated by inhibition of miR-199a-5p, and change in CHOP and caspase-12 expression induced by DSS was downregulated by inhibition of miR-199a-5p. \**P*< 0.05 \*\**P*< 0.01 vs. CON; ##*P*< 0.01 vs. DSS; ^^*P*< 0.01 vs. NC

## Discussion

miRNAs are a class of highly conserved, very short, and endogenous non-coding single-stranded RNAs that are about 22 nucleotides long; they were first discovered in eukaryotic nematodes in 1993 [7]. Recent studies [8, 9] have found that miRNA expression is specific in colonic mucosal tissues and is involved in various physiological and pathological processes of the development of UC. Notably, miR-199 is one of the most common and conserved miRNAs that has a variety of functions in vivo and plays an important role in cell replication [10], autophagy [11], and other functions. At present, most studies are on the mechanism of miR-199a-5p in digestive tract tumors and there are a few on UC.

Previous studies have shown [10] that miR-199a-5p is highly expressed in peripheral blood of patients with UC, so we believe that miR-199a-5p plays an important role in the pathological process of UC. In our study, a DSS-induced mouse model was established to verify the specific expression of miR-199a-5p and ERS marker protein XBP1 in colon mucosal tissues. Induced ERS by TG in HT-29 and differential expression of miR-199a-5p when ERS occurred were found in our study. Further, inhibition of miR-199a-5p can alleviate ERS and the associated apoptosis. Bioinformatics analysis showed that XBP1 might be a potential target of miR-199a-5p, and a luciferase reporter assay showed that miR-199a-5p inhibited its expression by directly binding to the 3′-UTR of XBP1. Consequently, a decreased expression of XBP1 can affect the ERS process. We obtained the above results by inhibiting the expression of miR-199a-5p in DSS-induced mouse model using an rAAV vector. Therefore, this study provides evidence for the first time that miR-199a-5p mediated ERS in the development of UC by regulating XBP1.

Previous studies have found that ERS can participate in the pathological process of UC by inducing cell apoptosis and playing a role in the innate immune response and epithelial autophagy [12, 13]. Impaired ERS eventually affects the secretion function of Pan’s cells and goblet cells; a decrease in Pan’s cells leads to abnormal function, reducing the intestinal resistance to microorganisms. Therefore, the functional stability of ERS-related proteins, as well as their regulation, is of great significance in understanding the occurrence and development of UC. The IRE1-XBP1 pathway is the most stable pathway in biological evolution and is activated in the early stage of ERS to promote cell survival. This pathway shuts down when a stimulus is strong and prolonged. Therefore, the open state of this pathway determines the ultimate condition of cells [14]. XBP1 can also negatively regulate apoptosis by affecting caspase-12, CHOP, and p-JNK [15], and its deletion can lead to the overactivation of IRE1 and trigger regulated Ire1-dependent decay (RIDD) in cells, resulting in apoptosis [14]. Therefore, this study implies that XBP1 plays an important regulatory role in the pathophysiological process of UC.

In this study, XBP1 expression was demonstrated to be decreased by miR-199a-5p. Inhibiting miR-199a-5p showed significant upregulation of XBP1 and remission of inflammation in DSS-induced colitis, which was consistent with previous findings that activation of XBP1 inhibited the pathogenesis of UC [16]. At present, there are limited studies on the regulation of XBP1 in UC. In our study, it was found that miR-199a-5p further affected the intracellular ERS and apoptosis processes by regulating XBP1. Therefore, in a DSS-induced mouse model, miR-199a-5p is upregulated in intestinal tissues, and changes corresponding to miR-199a-5p are also observed in XBP1-related functions, thereby affecting the ERS process. This provides a new theoretical basis for understanding the pathophysiological mechanism of UC and new therapeutic targets for drugs.

In conclusion, miR-199a-5p was upregulated by DSS treatment in mice and promoted ERS and cell apoptosis by targeting the 3′-UTR of XBP1, which is a key component of ER stress. Inhibition of miR-199a-5p prevented DSS-induced UC and cell apoptosis and increased XBP1 expression, suggesting that miR-199a-5p may be a potential diagnostic biomarker of UC and that XBP1 could be a potential therapeutic target. However, all data in the present study are preclinical results, so further clinical study is warranted to elucidate the effects of miR-199a-5p/XBP1 network in UC.

## Materials and Methods

### Cell culture and miRNA transfection

The human intestinal epithelial cell line, HT-29, was purchased from the American Type Culture Collection (Manassas, VA, USA). The cells were cultured with Dulbecco’s Modified Eagle Medium (DMEM), which was supplemented with heat-inactivated fetal bovine serum (FBS; Gibco, USA), penicillin (Gibco, USA), and streptomycin (Gibco, USA); they were incubated at 5% CO2 at 37 ° C.

We investigated whether Thapsigargin (TG) (American Sigma) can induce the intracellular ERS process [17], and the cells were divided into six groups: control group, TG group, TG + miR-199a-5p mimic group, TG + Negative Control (NC) mimic group, TG + miR-199a-5p inhibitor group, and TG + NC inhibitor group. Ten microliters of miR-199a-5p mimic or its negative control and miR-199a-5p inhibitor or its negative control (Genepharma, China) were dissolved in 100 μL opti-MEM(Gibco, USA). Then 100 μ L of this solution was gently mixed with 8 μ L of Lipofectamine^™^ 2000 (lipo) (Invitrogen, USA). A solution of lipo and siRNA (total volume 200 μ L) was prepared and incubated at room temperature for 15–20 min to form the complex, as per the manufacturer’ s instructions. HT-29 cells seeded in sterile 6-well plates were supplemented with 200 μ L of the mixture per well. The cells were cultured in a CO2 incubator at 37 ° C for 6 h, and the supernatant was replaced with complete MC5A (Gibco, USA) medium, followed by the addition of TG (1 mol/L) and incubation for 24 h. Each experiment was performed in triplicate.

### Animal model

We obtained 6- to 8-week-old adult male C57BL/6 mice from Beijing HFK Bioscience Co., Ltd. (Beijing, China). All mice were housed in a room with controlled temperature (21 ± 2 °C) and humidity (50 ± 5%), as well as a 12-h light/dark cycle. The mice had access to a standard mouse diet and sterile water ad libitum and were acclimated to the laboratory conditions for seven days before undergoing any experimental procedures. Recombinant adeno-associated virus (rAAV), which serve as vectors for inhibit miR-199a-5p and its NC (1.2*10^12vg/mL, Hanbio, China) were divided into 200-μL aliquots. Mice were assigned into 4 groups (n = 5 mice/group) with the following treatments: group I (Control), mice administered with normal drinking water; group II (DSS), mice administered with drinking water containing 3% (w/v) of dextran sulphate sodium (DSS) (160110; MP Biomedicals, Santa Ana, CA, USA) for one week after three weeks of normal drinking; group III (DSS + rAAV-inhibited miR-199a-5p), mice were administered with 200 μ L of rAAV-inhibited miR-199a-5p by enema on the first day and then administered with drinking water containing 3% of DSS for one week after three weeks of normal drinking; group IV (DSS + rAAV-NC), mice were administered with 200 μ L of rAAV-NC by enema on the first day and then administered with drinking water containing 3% of DSS for one week after three weeks of normal drinking. Starting from day 22 of the experiment, the dietary amount, weight, fecal matter characteristics, and activity of mice in each group were observed and recorded daily. Disease activity index (DAI) was calculated as follows: DAI = (weight loss score + fecal characteristics score + bleeding score)/3 [18]. All mice were sacrificed via cervical dislocation on day 29 and their colon tissues harvested for subsequent analyses. All experimental procedures were conducted in accordance with the institutional guidelines for the care and use of laboratory animals of Renmin Hospital at Wuhan University (Wuhan, China) and conformed to the regulations outlined in the National Institutes of Health Guide for Care and Use of Laboratory Animals. Experiments were performed under a project license (NO.: WDRM20191008) granted by Renmin hospital of Wuhan university.

### Histopathological analyses

A portion of the distal colon, which was fixed with 10% buffered formalin for over 24 h, was mounted in paraffin and stained with hematoxylin and eosin (H&E) before observation under a light microscope (Olympus BX53). The histological scoring system (Dieleman et al., 1998) was used to grade the severity of the tissue damage induced by DSS.

### RNA extraction and quantitative real-time PCR (qRT-PCR) assay

TRIzol reagent (Aidlab, China) was used to extract total RNA. Reverse transcription was achieved using the HiScript Reverse Transcriptase (VAZYME, China) according to the manufacturer’s instructions. Relative RNA expression was examined by qRT-PCR using a SYBR Green Master Mix (VAZYME, China) on the ABI QuantStudio 6 system (Applied Biosystems, Thermo Fisher Scientific, Foster City, USA). Subsequently, miRNA expression was normalized against U6 expression, and mRNA expression of each target gene was normalized against GAPDH expression. The data were calculated by means of the 2-ΔΔCt method. The primer sequences used in this study are shown in Table 1.

**Table 1:**
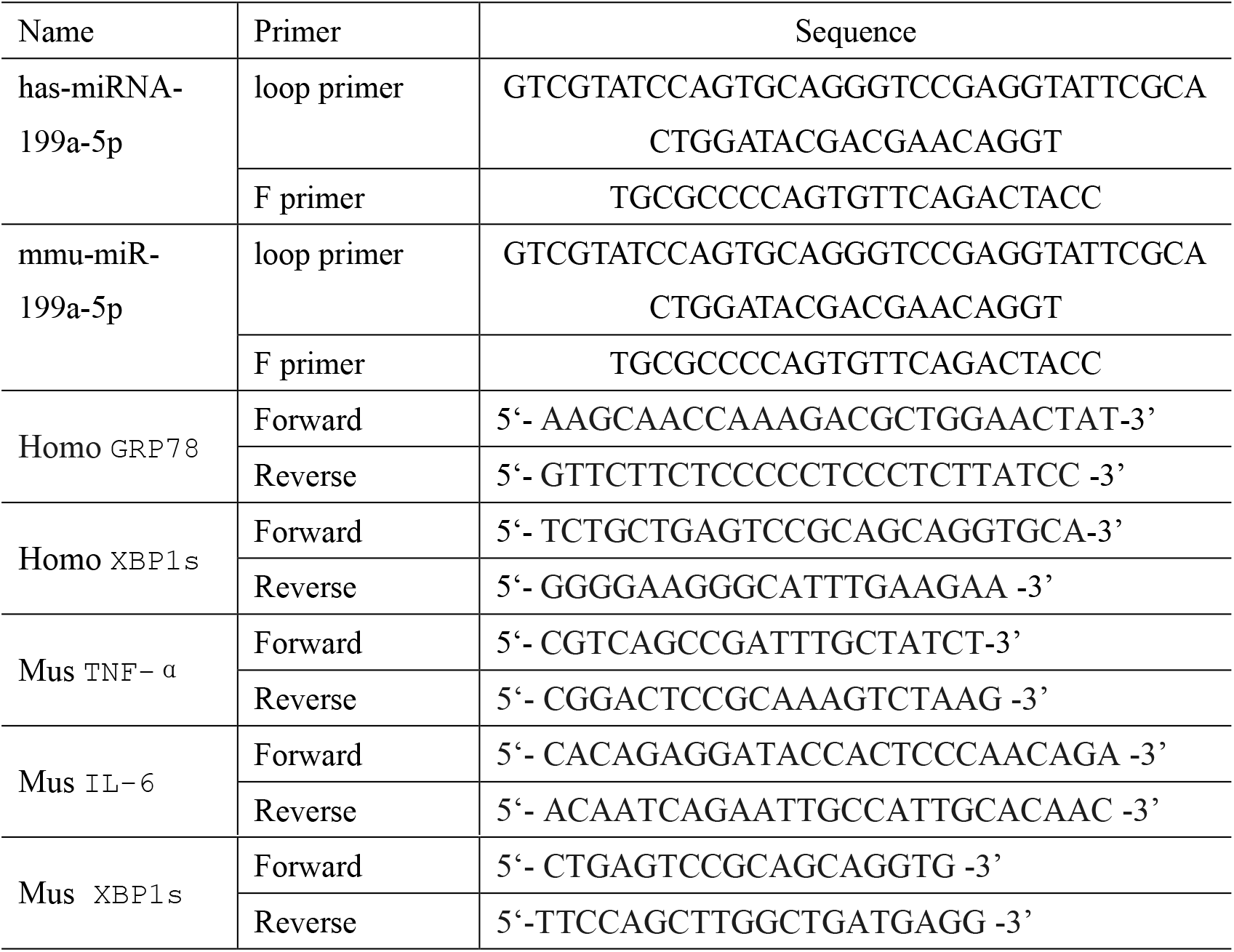
Primers used for qRT-PCR

### Western blotting

Protein lysates from colonic tissue samples were extracted using RIPA cell lysis buffer (Beyotime, China) and protein concentration was measured using BCA protein assay kit (Beyotime, China). Total protein (40 μ g) was electrophoresed on a 10% polyacrylamide gel by SDS-PAGE and transferred to PVDF membranes (Millipore, USA), followed by blocking with 5% milk for 2 h. Membranes were sequentially incubated with the following primary antibodies: GRP78(1: 1500 dilution, Proteintech), XBP1s (1:1000 dilution, Proteintech), Caspase-12 (1:1000 dilution, Abcam), CHOP (1:800 dilution, Proteintech) and GAPDH (1:1000 dilution, Goodhere). After incubation with primary antibodies overnight at 4 °C, the membranes were incubated once more with the appropriate secondary antibodies (Boster, China). Finally, the bands were detected using SignalFire^™^ ECL Reagent (Cell Signaling Technology, Inc., Danvers, MA, USA) and quantified using ImageJ software (NIH, Bethesda, MD).

### Luciferase reporter assay

Synthesized XBP1 3′ untranslated region (3′-UTR) that contained the predicted binding sites and mutated sequence were used. The 3′-UTR of XBP1 containing wide type (WT) and scrambled (MUT) miR-199a-5p binding sequence were inserted downstream of the firefly luciferase gene in pYr-MirTarget (YRGene, China) to generate the pYr-MirTarget-XBP1 3′UTR-WT and pYr-MirTarget-XBP1 3′UTR-MUT plasmid, respectively. The constructed plasmids were co-transfected into HEK293T cells with NC mimic or miR-199a-5p mimic using lipo. After 24 h, luciferase activity was assayed using the Luciferase Reporter Assay kit (Beyotime, China) according to the manufacturer’s protocol.

### Flow cytometry and terminal deoxynucleotidyl transferase dUTP nick end labeling (TUNEL)

Flow cytometry was used to detect HT-29 cell apoptosis using an AnnexinV-FITC/PI apoptosis detection kit (NanJing KeyGen Biotech Co.,Ltd.) according to the manufacturer’s protocol. The stained samples were analyzed with CytoFLEX (Beckman Coulter, Brea, CA, USA). The data on apoptotic cells are shown as two parameter dot-plots. Apoptotic DNA fragmentation was examined using a TUNEL Apoptosis Detection Kit (Roche Applied Science) according to the manufacturer’s protocol. Briefly, cells were plated in 24-well flat-bottom plates and fixed in 4% paraformaldehyde at 4 ° C for 25 min, permeabilized with 0.1% Triton X-100, and labeled with fluorescein-dUTP using terminal deoxynucleotidyl transferase. The localized red fluorescence of the apoptotic cells from the fluorescein-dUTP was detected via fluorescence microscopy (Olympus BX53).

### Statistical analysis

The data obtained from the experiments described above were analyzed using SPSS 23.0 software (IBM SPSS Inc., Armonk, NY, USA) and presented as mean ± standard deviation (SD). Differences between groups were assessed using one-way analysis of variance (ANOVA) and Student’s t-test. P values < 0.05 were considered to be statistically significant.

## Acknowledgements

Drs. Hesheng Luo confirmed that all authors have contributed to and agreed on the content of the manuscript.

## Author Contributions

Shanshan Wang, Lei Shen, Shuai Peng, Minxiu Tian and Xiangjie Li performed the experiments; Shanshan Wang and Lei Shen analyzed the data, prepared the figures and drafted the manuscript; Hesheng Luo edited and revised the manuscript; Shanshan Wang, Lei Shen and Hesheng Luo are responsible for conception and design of the research.

## Financial disclosure

Drs. Wang, Shen, Peng, Tian, Li and Luo and have no conflicts of interest or financial ties to disclose.

